# Algal growth and morphogenesis-promoting factors released by cold-adapted bacteria contribute to the resilience and morphogenesis of the seaweed *Ulva* (Chlorophyta) in Antarctica (Potter Cove)

**DOI:** 10.1101/2024.07.05.601910

**Authors:** Fatemeh Ghaderiardakani, Johann F. Ulrich, Emanuel Barth, Maria Liliana Quartino, Thomas Wichard

## Abstract

Macroalgae are found in a variety of marine vegetation ecosystems around the world, contributing significantly to global net primary production. In particular, the sea lettuce species, i.e., members of the genus *Ulva* (Chlorophyta), are located in many ecological niches and are characterized by excellent adaptability to environmental changes but depend on essential associated bacteria, which release algal growth and morphogenesis-promoting-factors (AGMPFs). Our work investigated the hypothesis that bacteria need to be stress-adapted to provide sufficient amounts of AGMPFs for the growth and morphogenesis of *Ulva* throughout its life cycle, even under severe environmental conditions.

Our study thus aimed to understand which bacteria contribute to overcoming a variety of stressors in polar regions. Green macroalgae were collected from Potter Cove, King George Island (Isla 25 de Mayo), Antarctica, to study the associated microbiome and, subsequently, to identify AGMPFs releasing bacteria. Therefore, microbiome analysis was combined with morphogenetic bioassays and chemical analysis, identifying bacteria essential for algal growth under Antarctic conditions. Hereby, axenic cultures of a Mediterranean *Ulva compressa* (cultivar *Ulva mutabilis*), previously developed as a model system for bacteria-induced algal growth and morphogenesis, were inoculated with freshly isolated and cultivable Antarctic bacteria to determine their morphogenetic activity.

The exploratory microbiome investigation identified numerous cold-adapted AGMPF-producing bacteria. Unlike the reference bacterial strains isolated from the Mediterranean Sea, the cold-adapted isolates *Maribacter* sp. BPC-D8 and *Sulfitobacter* sp. BPC-C4, released sufficient amounts of AGMPFs, such as thallusin, necessary for algal morphogenesis even at 2°C. Our results illustrate the role of chemical mediators provided by bacteria in cross-kingdom interactions under cold conditions within aquatic systems. The newly isolated bacteria will enable further functional studies to understand the resilience of the holobiont *Ulva* and might applied in algal aquaculture even under adverse conditions. The study highlights the importance of ecophysiological assays in microbiome analysis.

## 1. Introduction

Seaweeds (marine macroalgae) are known as “ecosystem engineers” due to their essential roles in global ecology (Jones et al. 1994). They are cosmopolitan and have become a source of valuable, sustainable biomass in the food, feed, chemical, and pharmaceutical industries (Charrier et al. 2017). Given the pivotal importance of macroalgae, a deep understanding of their biology and physiology—particularly in terms of their development, morphogenesis, life cycle regulation, and evolutionary strategies—will facilitate the engineering of new and valuable biological systems, improvements in the productivity of seaweed cultivation systems, and reduction of the deleterious ecological, environmental, and economic effects of macroalgae (e.g., nuisance blooms and biofouling).

Sea lettuce (belonging to the genus *Ulva* of Chlorophyta) is characterized by its adaptability to environmental changes. It is often found in waters heavily polluted by humans and under extreme temperatures, such as in polar or tropical regions. To survive as the currently established population, macroalgae must have successfully resisted a multitude of stressors, such as cold temperatures, dramatic changes in light, and large doses of UV radiation. To understand the effects of these factors, including global warming, on biodiversity or ecosystem features, a greater understanding of the stress mitigation techniques and adaptive processes of these species is needed.

The close relationship between macroalgae and bacteria, particularly in extreme ecosystems such as that of Antarctica, represents an exceptional study system for better understanding host–microbiome adaptation to harsh environmental conditions, the associated adaptive mechanisms, and the specific contribution of each compartment involved in algal–bacterial interactions to the functioning of the Antarctic marine ecosystem (Ghaderiardakani et al. 2022).

The functional importance of associated microbial communities in other terrestrial ecosystems (mainly plants) and the development of host resilience to various stresses have been the focus of many investigations (Chojak-Koźniewska et al. 2018; de Vries et al. 2020). Similarly, in many ways, marine life depends on symbiotic interactions with microbes. For instance, microbial communities control various aspects of the biology and ecology of algal hosts, including growth, morphogenesis (Wichard 2015), adaptation, acclimation (Dittami et al. 2016; Ghaderiardakani et al. 2020), resilience, health (Clements and Hay 2023), reproductive performance and settlement (Callow and Callow 2006). In turn, algae represent a rich source of organic components recognized as potential primary chemical signals that drive the dynamics of bacterial communities within the algal–bacterial alliance (Seymour et al. 2017; Kessler et al. 2018; Alsufyani et al. 2020).

Sampling a holobiont (i.e., an alga and its associated bacteria) from its natural but harsh environment is one of the possible experimental approaches for identifying adaptation strategies. For example, marine macroalgae in the southern polar region experience very low water temperatures (−1.8 °C in winter and a maximum of 5 °C in the summer) throughout the year (Wiencke and Dieck 1990; Eggert and Wiencke 2000). When considering the strategies of macroalgae for adaptation to the lowest temperatures, their associated bacterial community must be considered to understand how life is maintained under such extreme conditions. For example, it has been suggested that carotenoid-producing bacteria could enable survival under the unique environmental conditions in Antarctica, such as the low temperature, drastic changes in light, freezing-thaw events, high doses of UV radiation, and relatively low humidity (Alvarado et al. 2018; Vila et al. 2019). Interestingly, despite the distribution of seaweeds across different parts of the vertical zone of the Antarctic coast, the symbiotic bacterial composition varies among different seaweed species. This variation highlights the potential (partial) role of algal hosts in determining the composition of epiphytic bacteria within the holobiont (Gaitan-Espitia and Schmid 2020). However, the structure, phylogenetic diversity, and functional role of the microbial communities associated with Antarctic macroalgae from the perspective of the chemical ecology of cross-kingdom interactions and the possible survival strategies involved in such stress-moderating interactions have not yet been well explored.

Many studies have investigated whether host dynamics, phylogeny, or ecology modulate the composition of microbial communities. In a study of the green macroalga *Ulva australis*, high phylogenetic variability in the associated microbial composition was identified, but interestingly, only a small amount of functional variability among the microbes was found, which suggests that a core set of functional genes is present within alga-associated microbial communities (Burke et al. 2011). Different microbial communities with similar functional characteristics can be associated with the same or even different algal species (Ghaderiardakani et al. 2017). Therefore, the existence of specific microbial functional genes rather than phylogenetic biofilm composition is the key factor shaping the microbial communities on the surface of the algal host. However, these functions may be broadly distributed across various taxonomic groups (Burke et al. 2011). In addition, functional characteristics within the same microbiome can also change in a changing environment (Hmani et al. 2023). By employing mass spectrometry profiling and imaging techniques, specific metabolic markers have been introduced to visualize the distribution of bacteria (the producers of metabolic markers) on the surface of *Ulva* spp. (Vallet et al. 2021).

One of the well-studied model systems for investigating the role of cross-kingdom interactions in the completion of morphogenesis and development is a tripartite model system including the axenic green alga *Ulva mutabilis* and its two associated essential bacteria (*Roseovarius* sp. strain MS2 and *Maribacter* sp. strain MS6) (Wichard 2015; Blomme et al. 2023). Without the bacteria, the gametes of *U. mutabilis* develop into a callus-like biomass characterized by slow growth, atypical cell wall formation with protrusions, and a lack of thalli and rhizoids (Spoerner et al. 2012; Grueneberg et al. 2016). However, within the tripartite system, *Roseovarius* sp. strain MS2, via a yet-unknown compound, induces cell division (Wichard 2023), and in turn, these morphogen-producing bacteria benefit from the interaction by using the glycerol released by *Ulva* as a primary carbon source (Kessler et al. 2018). The second essential bacterium, *Maribacter* sp. strain MS6, triggers cell differentiation, cell wall development, and rhizoid formation in *U. mutabilis* by releasing the bacterial morphogen thallusin, the only identified algal growth and morphogenesis-promoting factor (AGMPF) in *Ulva* (Wichard et al. 2015; Alsufyani et al. 2020). At very low concentrations (EC_50_ = 5 pmol L^−1^), the morphogen thallusin induces rhizoid and cell wall formation (Alsufyani et al. 2020; Dhiman et al. 2022; Ulrich et al. 2022). Thallusin was initially isolated from the bacteria *Cytophaga* sp. and *Zobellia* sp., which promote thallus development in *Monostroma oxyspermum* (Matsuo et al. 2003; Matsuo et al. 2005). In fact, thallusin functions as a chemical mediator of algal development. Like plant hormones, thallusin has distinct functions in algal development depending on the recipient species and works synergistically with unknown *Roseovarius*-released factors as an inducer of algal morphogenesis in *Ulva* spp. (Alsufyani et al. 2020). The model organisms *U. mutabilis* and *U. compressa* were previously designated as being conspecific (Steinhagen et al., 2019). We will refer to *U. mutabilis* in this study to be consistent with the literature and avoid confusion with prior studies in which natural isolates were predominantly characterized based on physical features.

Since the bacterial impacts on the production of morphogenic substances occur at very low concentrations, detailed mechanistic studies are necessary to determine possible changes in the microbiome. To elucidate the complex interactions in the holobiont of *Ulva* and to propel the exploration of adaptation mechanisms, we developed a holistic top-down approach beginning with microbiome analysis to identify cold-adapted bacteria that release potentially AGMPFs under adverse conditions via morphogenetic *Ulva* bioassays followed by the quantification of thallusin production in bacterial media exposed to cold temperatures. We hypothesized that *Ulva* could respond adequately to stress if its microbiome adapted to environmental changes to provide the necessary algal growth-promoting factors.

## 2. Methods

### 2.1 Collection site and sampling of macroalgae and bacteria

The sampling locations were selected in the ice-free region in Potter Cove and Potter Peninsula (Isla 25 de Mayo/King George Island), Antarctica (62°14’S 58° 31’W) (**Fig. 1A**) (Wiencke et al. 1998; Quartino et al. 2005; Fromm et al. 2020). Approximately 50 algal specimens of green algae, such as *Ulva* spp. and *Monostroma* spp., were collected from the Antarctic Specially Protected Area (ASPA 132), and water samples were collected from the coastline at low tide in January/February 2020 (**Fig. 1B**). The samples were transported to the Dallmann Laboratory operated by the Alfred Wegener Institute (AWI) and Instituto Antártico Argentino (IAA) adjacent to the Argentinean Carlini Station.

**Figure 1.**
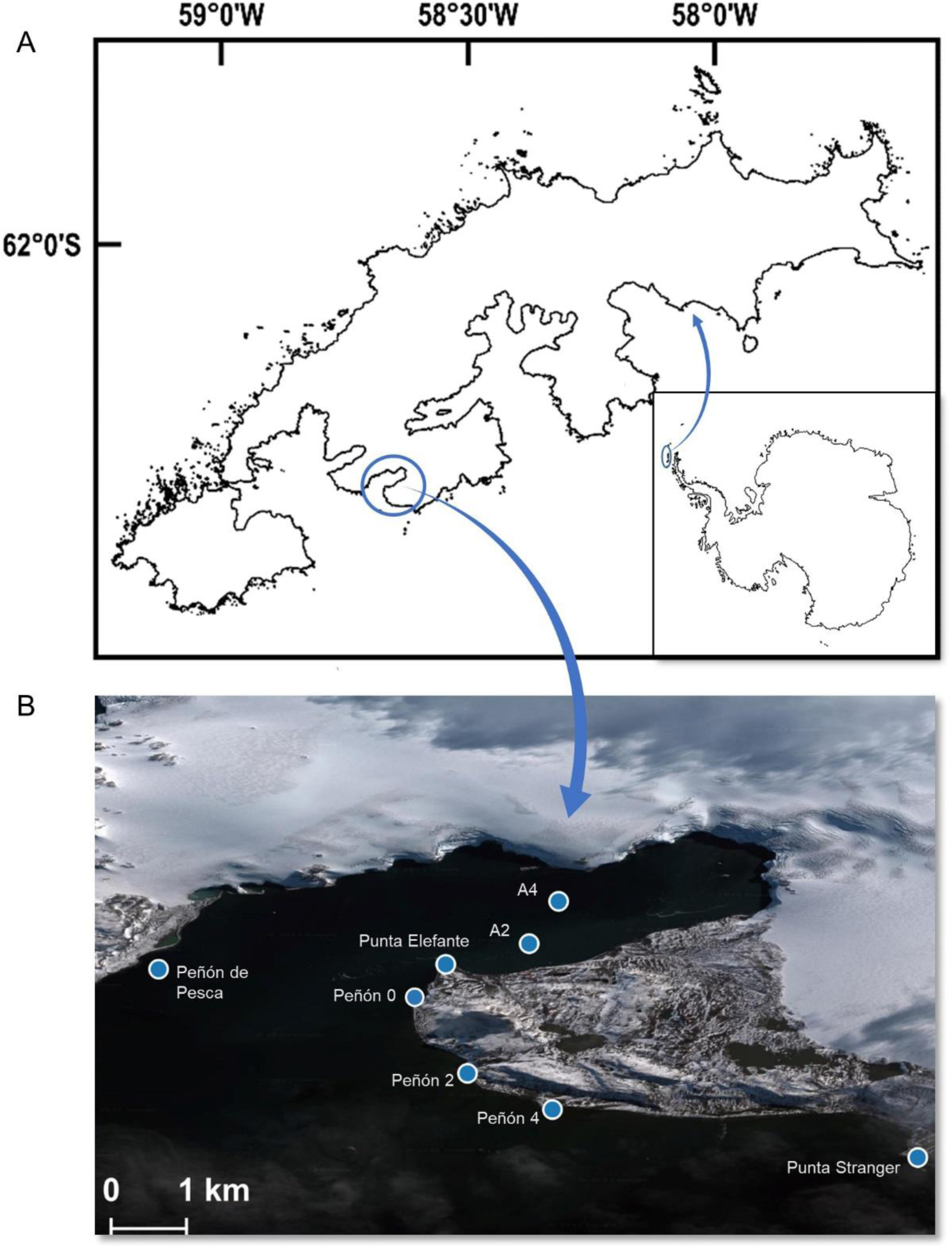
Sampling sites. (**A**) Location of the studied sites on Potter Cove and Potter Peninsula on Isla 25 de Mayo/King George Island within the Shetland Archipelago, Antarctica (Potter Cove, Google Maps, 1 January 2023). (**B**) Eight sampling sites are shown along the coastline and in Potter Cove, including Peñón de Pesca (PDP), Punta Elefante (PE), and Punta Stranger (PS).

### 2.2 Taxonomic classification of the macroalgae collected in Potter Cove

The macroalgal specimens were collected from Potter Cove, near the Carlini Station and the intertidal platform further outside the bay on King George Island, to determine the green macroalgal abundance, species composition, and community structure in the habitat (**Fig. 2**). The samples were frozen and maintained at −80 °C during shipment to the Alfred Wegener Institute (Bremerhaven, Germany).

**Figure 2.**
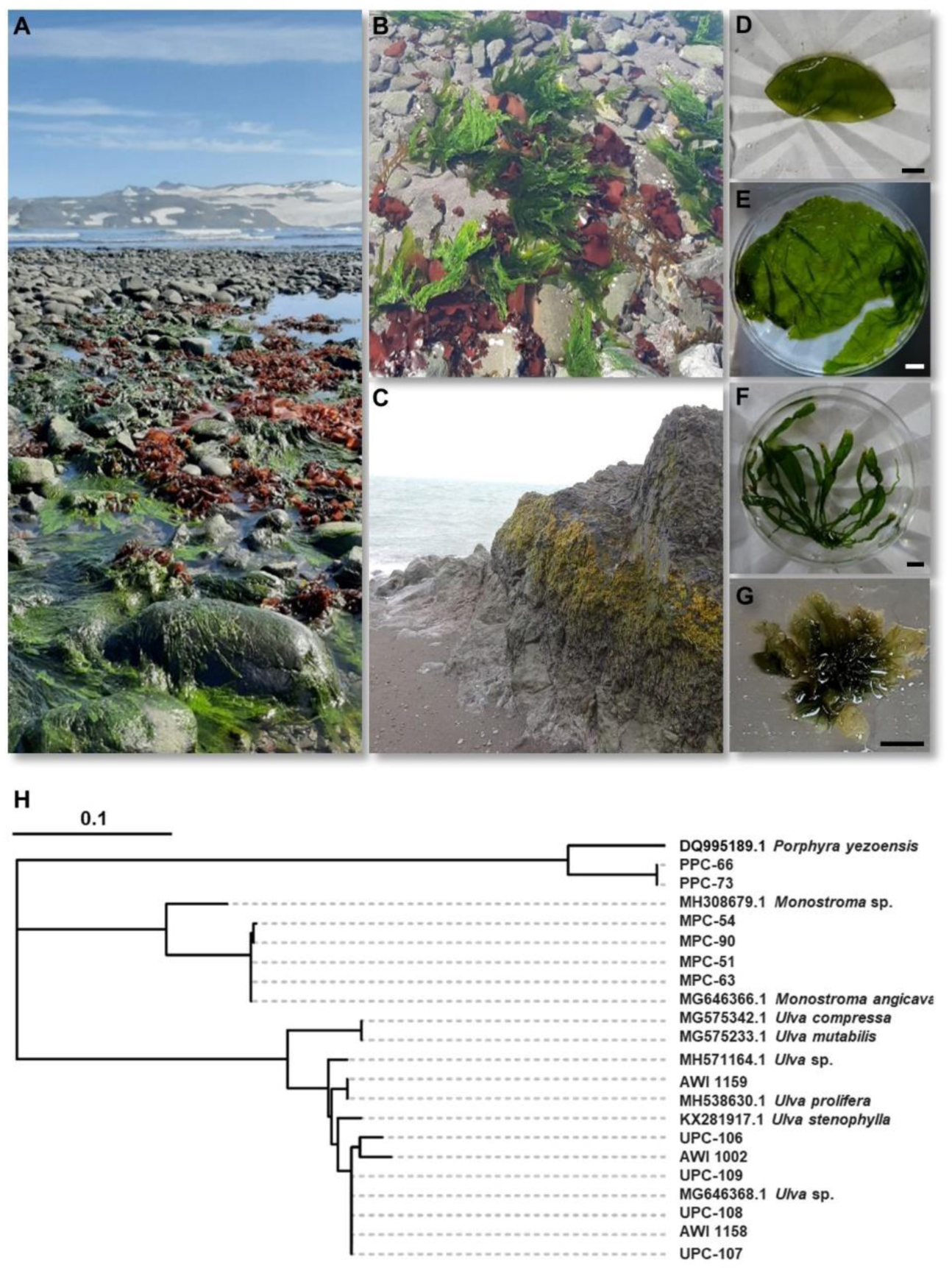
Diversity of green macroalgae sampled in Potter Cove of King George Island (Antarctica). (**A**) The diversity and abundance of various macroalgae (green, red, and brown) are illustrated in this photograph. (**B-F**) The figures demonstrate that the green algal samples were predominantly distributed in the intertidal zone. Scale bars = 1 cm. (**A-D**) were taken with permission from Fromm et al., 2020. (**G**) A phylogenetic tree was inferred from *tuf*A sequences and constructed with the maximum likelihood method (MEGA 10.1.8; Tamura et al., 2011). The Mediterranean model organism *U. mutabilis* Føyn (Føyn 1958), belonging to *"Ulva-compressa-pseudocuvata"* (Steinhagen et al. 2019), was used as an internal reference, and *Porphyra yezoensis* was used as an outgroup.

For genotyping, a tissue sample (up to 100 mg fresh weight per macroalga) was used for genomic DNA extraction using the GenElute Plant Genomic DNA Miniprep Kit (Merck). Genotyping PCR was performed using Prime-STAR GXL DNA Polymerase (Takara) in the presence of 5% DMSO (Kwantes and Wichard 2022) with the primer pair designed for tufA (forward: GGNGCNGCNCAAATGGAYGG, reverse: CCTTCNCGAATMGCRAAWCGC) (Famà et al. 2002).

The biosamples were named according to their identification details, such as organism type, location, and identifier, and were archived at the National Center for Biotechnology Information (NCBI; Supplementary Material: Table S1).

### 2.3 Metagenomic analysis of surface-associated bacteria

After systematically checking for sessile organisms, which, if present, were carefully removed with a sterile scraper, the algal tissue was rinsed three times in autoclaved *Ulva* culture medium to remove unattached bacteria from the surface. Three replicates of 200 mg of algal tissue were cut into small pieces, rapidly frozen with liquid nitrogen, and stored at −80 °C until DNA extraction. According to the manufacturer’s protocol, microbial metagenomic DNA was extracted using the DNeasy® Power Soil® Kit (Qiagen, Germany). The primers 515F-Y "new EMP" GTGYCAGCMGCCGCGGTAA and 806RB "new EMP" GGACTACNVGGGTWTCTAAT, which target the V4 region of the 16S rRNA gene, were used for sequencing via the Illumina MiSeq 3 platform. Microbiome analysis was initiated by performing quality control and investigating the raw sequencing reads using FastQC v0.11.9 (http://www.bioinformatics.babraham.ac.uk/projects/fastqc). Quality filtering and trimming were performed using prinseq (version 1.2.4) with the following parameters: -min_qual_mean 20 -ns_max_n 0 -noiupac -trim_qual_right (Schmieder and Edwards 2011). The DADA2 pipeline v1.20.0 (Callahan et al. 2016) in R v4.0.3 was used to process the resulting reads and infer exact amplicon sequence variants (ASVs) from the paired-end reads. The forward and reverse reads were merged to obtain fragments ranging from 220 to 263 nucleotides in length, with 93.8% having a length of 253 nucleotides. The sequences were grouped by 100% similarity to generate unique ASVs, and taxonomic identities were assigned to them using the pretrained naïve Bayes classifier on the DADA2-formatted SILVA database (silva_nr99_v. 138.1) (Quast et al. 2013) (**Fig. 3**; NCBI accession numbers of the raw sequencing data: PRJNA828511, SAMN39202330-SAMN39202384; Supplementary Material: Table S2).

**Figure 3.**
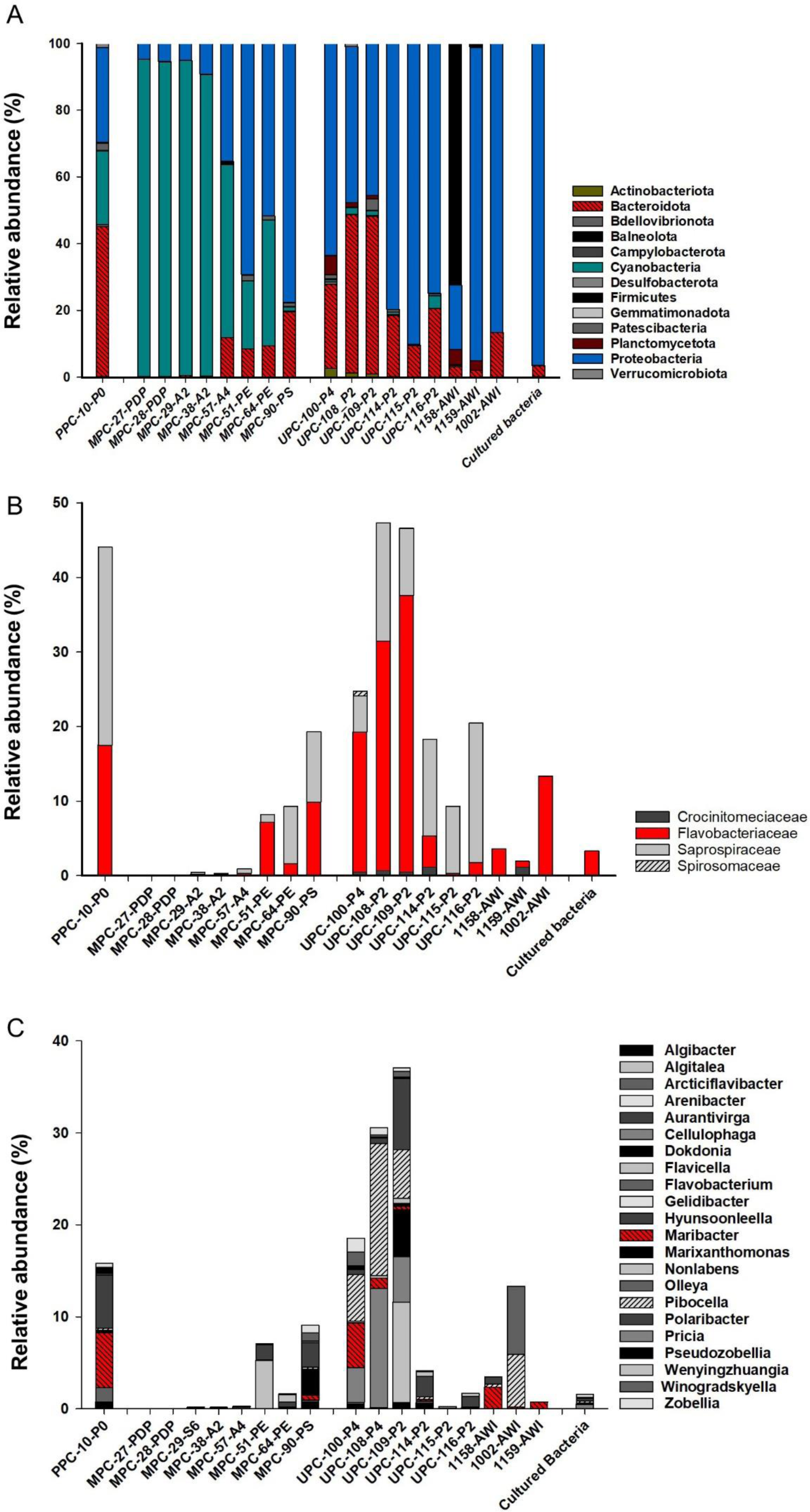
Relative abundance and prevalence of surface microbes on green macroalgae collected across all the sampling sites of Potter Cove (King George Island, Antarctica). The diversity of the microbiota was compared with that of a mixed sample of bacteria cultivated in marine broth. (**A**) The relative abundance of whole phyla in the microbiome. (**B**) The relative abundance of bacteria in the Bacteroidota family. (**C**) The taxonomic profile of the genera assigned to the family Flavobacteriaceae. (PPC, UPC, MPC: *Porphyra*, *Ulva* or *Monostroma* collected from Potter Cove; AWI: Antarctic *Ulva* sample from the Alfred-Wegener-Institute culture collection). The average of three measurements is shown (*n* = 3). A mixture of isolated and subsequent “cultured bacteria” was also analyzed (*n* = 4). Species names are arranged alphabetically from top to bottom (*Zobellia* - *Algibacter*). The hatched bars indicate the genera *Maribacter* and *Pibocella*, which were of particular interest in this study.

### 2.4 Identification of bacteria that release essential AGMPFs using the standardized model system of *Ulva mutabilis*

Microbiome analysis was accompanied by functional examinations to investigate growth and morphogenesis-promoting traits within the microbiome (Hmani et al. 2023). For this, bacteria were isolated from the surface of the algal samples using sterile swabs. They were streaked onto preprepared marine agar (MA) plates (Roth, Karlsruhe, Germany, marine broth (MB) supplemented with 1% agar) to obtain single colonies (Grueneberg et al. 2016). All the plates were incubated at 8 °C for three days. After 1–7 days, 443 distinct colonies were selected and transferred with sterile toothpicks into 96-well plates containing 200 μL of MA in each well.

To evaluate the activity of potential morphogenesis-inducing bacteria, a morphogenetic bioassay was performed using axenic cultures of *U. mutabilis* (Grueneberg et al. 2016). Axenic gametes of *U. mutabilis* (strain FSU-UM5-1) were prepared according to the method established by Califano and Wichard (2018) and inoculated with bacteria isolated from green or red macroalgae collected individually from Potter Cove, Antarctica (section 2.1). As positive controls, the AGMPF-releasing reference strains *Roseovarius* sp. (GenBank EU359909) and *Maribacter* sp. (GenBank EU359911) were inoculated singly and jointly with the axenic cultures (Spoerner et al. 2012; Ghaderiardakani et al. 2017). As a negative control, some wells were randomly retained without bacterial inoculation (only axenic gametes) in each row (**Fig. 4**). All inoculations and controls were carried out in triplicate. After inoculation, the final optical density of the bacteria was OD_620_ = 1.0 × 10^−4,^ and all the cultures were incubated at 18 °C.

**Figure 4.**
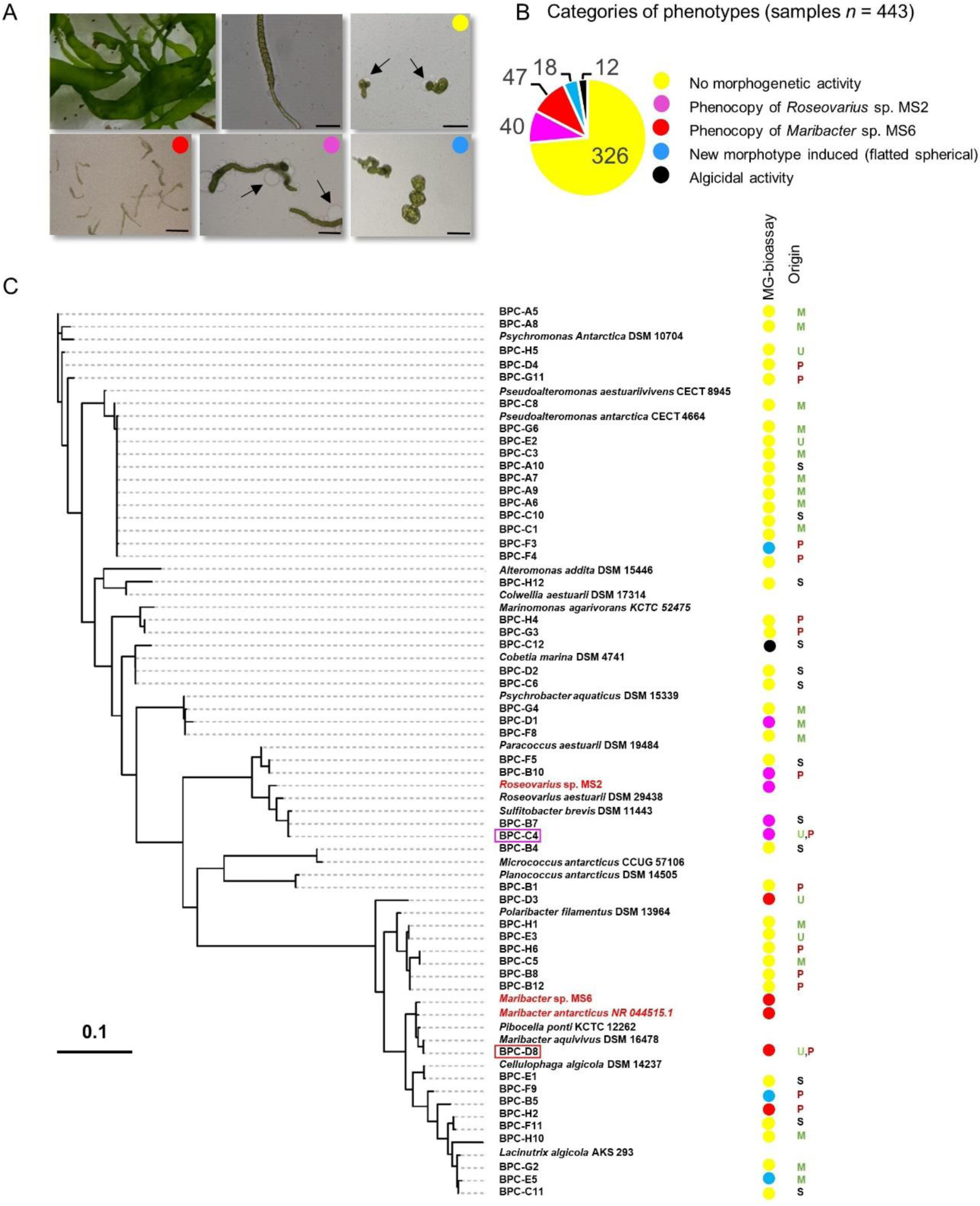
Morphogenetic bioassays with *Ulva mutabilis* and identification of isolated bacteria across all the sampling sites in Potter Cove (King George Island, Antarctica). (**A**) The figure exemplifies purified *Ulva mutabilis* germlings inoculated with bacteria isolated from green macroalgae collected in Potter Cove (Antarctica). Scale bars = 100 µm. (**B**) In total, 443 bacterial isolates from Potter Cove were tested and categorized. (**C**) The phylogenetic tree is based on 50 selected bacteria and their 16S rRNA gene sequences. *Roseovarius* sp. MS2, *Maribacter* sp. MS6 and *Maribacter antarcticus* DSM 21422 were used as reference strains. The results of the morphogenetic (MG) bioassay and the origin of the bacteria are annotated (M: *Monostroma* spp.; U: *Ulva* spp.; P: *Porphyra*; S: surface seawater).

### 2.5 Molecular identification and phylogenetic analysis of the cultivable bacteria collected from the algal surface

DNA was extracted from 49 out of 443 cultivable cold-adapted bacterial isolates according to the manufacturer’s instructions for the DNeasy Blood and Tissue Kit (Qiagen, Germany) (Grueneberg et al. 2016). To identify the bacteria, partial 16S rDNA sequences (∼1500 bp) were amplified from these strains using the primer pair 27f (GGG TTT GAT CCT GGC TCA G) and 1390r (ACG GGC GGT GTG TRC AA) (Frank et al. 2008). The preparation of the reaction mixture and amplification of 16S rRNA were performed following the protocol described by Ghaderiardakani et al. (2017). The PCR products were then subjected to forward primer-mediated sequencing using the chain termination method (Eurofins, Germany). The closest homologous sequences in the GenBank database were identified and used to construct the phylogenetic tree.

Taxonomic assignment of the isolated and cultivated bacteria was performed using the Ribosomal Database Project classifier tool (RDP, release 10, http://rdp.cme.msu.edu) at an 80% confidence threshold. Bacterial-type strains representing the closest 16*S* rRNA gene relatives to the sequences generated in this study were identified using the RDP sequence match tool and were included in the phylogenetic inference efforts for taxonomic reliability. For phylogenetic reconstruction, 16*S* rRNA gene sequences from our isolates, their closest type strains, and reference *Ulva*-associated strains were aligned by the SINA web aligner tool within the ARB-SILVA database (**Fig. 4b**; NCBI accession numbers: OR957431-OR957479; Supplementary Material: Table S3).

### 2.6 Bacterial whole-genome-based phylogeny

The bacterial isolate BPC-D8, which was obtained from the algal surface and tested positive for morphogenetic activity, was cultivated for 4 days in MB medium in sterile polycarbonate flasks (Sarstedt, Germany) at 20 ± 1 °C in natural light with orbital shaking. Genomic DNA was extracted using the cetyltrimethylammonium bromide protocol (William et al., 2012). The DNA sample was sheared in Covaris g-TUBEs, resulting in fragments with a target size of 20 kb. A Pacific Biosciences (PacBio) SMRTbell Express library was prepared according to the manufacturer’s instructions using AMPure PB beads for size selection. Microsynth AG (Switzerland) performed library barcoding and pooling/multiplexing, library quality control (Femto Pulse analysis and BluePippin size selection), genome sequencing on a PacBio Sequel sequencer, and subsequent assembly. After subread generation (PacBio single-molecule real-time [SMRT] Link v.10.1) and extraction (SAMtools v.1.14), Filtlong v.0.2.1 was used to filter subreads smaller than 1,000 bp. *De novo* assembly was performed using the Flye assembler (v. 2.8.3). The default parameters were applied for all the software packages. Error correction and Flye software enabled the detection of circularity. The taxonomic classification was determined using the Genome Taxonomy Database (GTDB) with GTDB-Tk v1.7.0. The taxonomic method was based on the classification defined by topology and average nucleotide identity (ANI)-based comparisons (https://www.kbase.us/) (Jain et al. 2018; Chaumeil et al. 2020) (**Fig. 5**).

**Figure 5.**
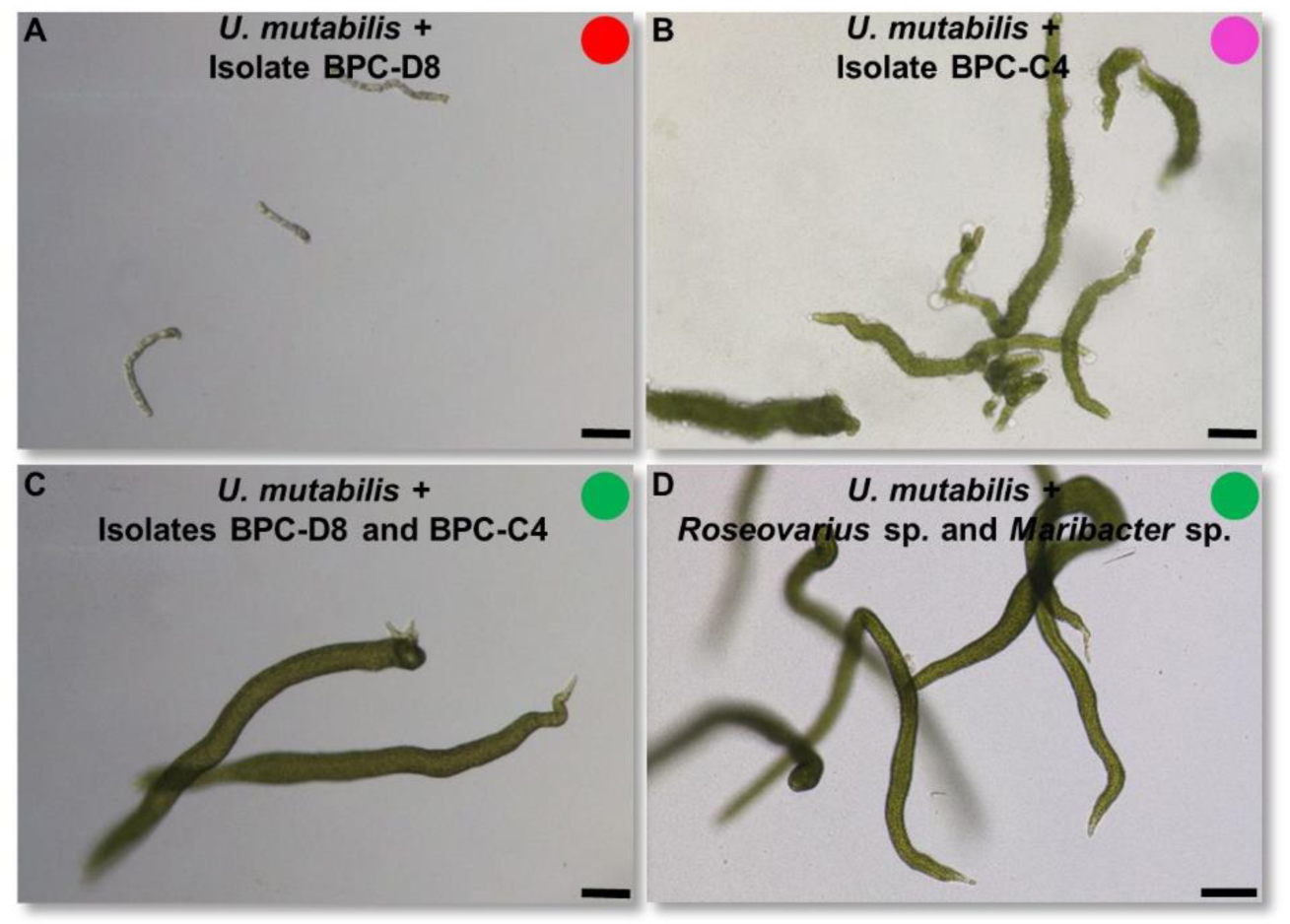
A combination of two Antarctic bacteria induces the complete morphogenesis of *Ulva mutabilis*. Comparison of the bacterium-induced morphogenesis of *U. mutabilis* through Antarctic bacteria (**A-C**) and Mediterranean bacteria (**D**) isolated by Spoerner et al. (2012). Scale bar = 100 µm.

### 2.7 Thallusin quantification in cold-adapted bacterial cultures

The bacteria were cultivated in 200 mL of HaHa_100 growth medium (Hahnke et al. 2015) on an orbital shaker (230 V Euro Plug, Standard 5000, VWR). The optical density (OD) of the cultures at a wavelength of 620 nm was measured daily using a UV‒Vis spectrophotometer (Genesys 10S UV‒Vis Spectrophotometer, Thermo Scientific, Waltham, MA, USA). A calibration curve of OD versus cell count was prepared to determine cell density (Ulrich et al. 2022). Thallusin quantification was performed on an ultrahigh-performance liquid chromatography (UHPLC) system linked to a Q Exactive Orbitrap mass spectrometer (Thermo Fisher Scientific). Thallusin was extracted using C_18_ solid phase extraction followed by a derivatization step with iodomethane to prevent the formation of Fe–thallusin complexes, which would interfere with chromatographic separation (Ulrich et al. 2022) (**Fig. 6**).

**Figure 6.**
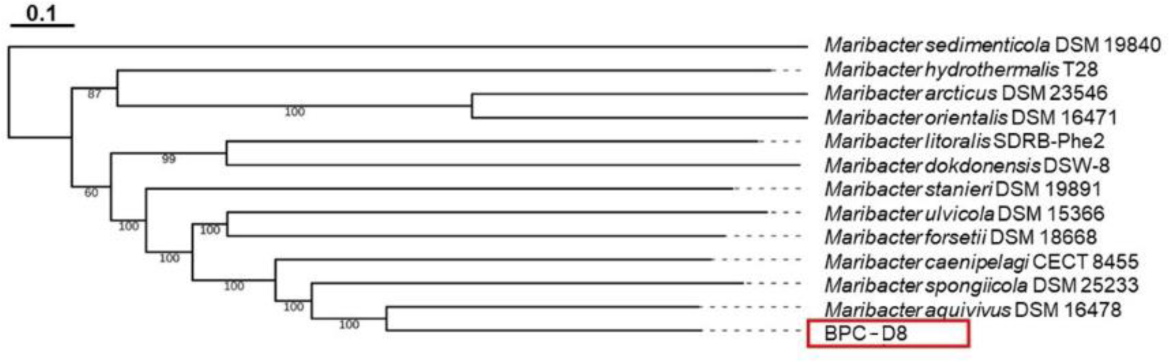
Phylogenetic analysis of the whole genome of the bacterial isolate BPC-D8. The strain was compared with the type strain genomes of publicly available genome sequences of *Maribacter* spp. The numbers beside each branch represent the bootstrap values from 100 replications.

## 3. Results and discussion

Given the worldwide distribution of green macroalgae, these organisms have managed to withstand a variety of stressors to become the established populations we are familiar with today. To understand the adaptations of *Ulva* to Antarctic conditions, macroalgae were collected from Potter Cove and Potter Peninsula, Isla 25 de Mayo/King George Island (**Fig. 1**, **Table 1**). The microbiome of the algal surface was assessed, and the bacteria-releasing AGMPFs were isolated. The production and prevalence of the AGMPF thallusin under cold stress was a particular criterion examined in the bacterial survey.

**Table 1:** Sampling site characteristics at Carlini Station, Potter Cove, Isla 25 de Mayo/King George Island.

| Location | Sample # | Latitude, Longitude | Date of sampling | Habitat | Collected macroalgae |
| --- | --- | --- | --- | --- | --- |
| **A2** | 29,38 | S 62° 14' 3.12"<br>W 58° 40' 3.00" | 01/02/2020 | Bay<br>(> 5 m depth) | *Monostroma* spp.<br>(floating) |
| **A4** | 57 | S 62° 13' 43.68"<br>W 58° 39' 49.32" | 04/02/2020 | Bay<br>(>5 m depth) | *Monostroma* spp.<br>(sessile) |
| **Punta Elefante (PE)** | 51, 54, 63, 64 | S 62°14'12.8"<br>W 58°40'41.5" | 04/02/2020 | Coastal water | *Monostroma* spp.<br>(sessile) |
| **Peñón 0 (P0)** | 10, 66, 73 |  | 29/01/2020 | Rocks | *Porphyra* spp.<br>(sessile) |
| **Peñón 2 (P2)** | 106,107,108,109,<br>114,115,116 | S 62°15'02.9"<br>W 58°40'31.4" | 09/02/2020 | Coastal water | *Ulva* spp.<br>(sessile) |
| **Peñón 4 (P4)** | 100 | S 62°15'19.4"<br>W 58°39'52.2" | 09/02/2020 | Coastal water | *Ulva* spp.<br>(sessile) |
| **Punta Stranger (PS)** | 90 | S 62° 15' 41.76"<br>W 58° 37' 4.08" | 08/02/2020 | Coastal water | *Monostroma* spp.<br>(floating) |
| **Peñón de Pesca (PDP)** | 27,28 | S 62°14' 16.5"<br>W 58° 42' 16.5" | 30/01/2020 | Bay<br>(>5 m depth) | *Monostroma* spp.<br>(floating) |

### 3.1 Collection and identification of Antarctic seaweeds in Potter Cove

In the Antarctic summer of 2020, the Antarctic coastline at Potter Cove was covered with brown, red, and, in some places, green macroalgae (**Fig. 2A-C**), whereas *Monostroma* spp. were found free-floating in water, and sessile *Ulva* spp. were only occasionally found along the coastline (**Figs. 1**, **2**). The red alga *Porphyra* sp. was identified at different locations, settled on rocks and stones (**Fig. 2C**).

The green macroalgae discovered were mostly monostromatic (**Fig. 2D, E**), mixed with a few distromatic foliose specimens (**Fig. 2F**). Green macroalgal samples (indexed as MPC: *Monostroma* Potter Cove and UPC: *Ulva* Potter Cove) were collected from all the designated sampling stations throughout our excursion, with an emphasis on the distromatic specimens (**Fig. 1**). Red algae were collected for comparison of the related microbiomes (PPC: *Porphyra* Potter Cove). DNA sequences of the gene *tufA*, encoding the elongation factor Tu, were determined from 10 macroalgal isolates and compared with available sequences in GenBank using the Basic Local Alignment Search Tool.

*TufA* sequences revealed that MPC isolates #54, #90, #51, and #63 were closely related to *Monostroma angicava* (**Fig. 2D, F**). In contrast, UPC isolates #106, #109, #108, and #107 were phylogenetically close to *Ulva* strains such as *Ulva linza* and *Ulva prolifera* (**Fig. 2H**). Moreover, *tufA* sequences of the collected green macroalgae aligned with those of strains isolated near the Great Wall Station on King George Island in 2014 (*Ulva* sp.: MG646368.1; *Monostroma* sp.: MG646366.1).

Interestingly, UPC-106 and UPC-108,107 were phylogenetically close to AWI isolates #1002 and #1158, which were collected 30 years ago in the same region (AWI #1002 was originally taxonomically attributed to *Ulva bulbosa*).

Approximately 120–130 macroalgal species have been recorded in Antarctica, mainly from the western Antarctic area along the peninsula (Wiencke and Clayton 2002). Interestingly, a relatively high proportion of marine flora endemism has been identified in this area (Wiencke and Clayton 2002). The green algae reported from the region between King George Island and Anvers Island include *Prasiola crispa* in the supralittoral zone and other species, such as *Urospora penicilliformis*, *Ulothrix* spp., *Enteromorpha bulbosa,* and *Acrosiphonia arcta*, which were found several times on rocks or in tidal pools approximately 30 years ago (Wiencke and Clayton 2002; Wiencke et al. 2007), but the species differed from the *Monostroma* and *Ulva* strains isolated in this study. The bacteria on the macroalgal surface of those isolates were analyzed to identify candidate algal symbionts releasing AGMPFs.

### 3.2 Microbiome analysis of macroalgae collected from Antarctica as a guide for the isolation of AGMPF-producing bacteria

To provide a comprehensive overview of the bacteria associated with green macroalgae, the surface microbiomes of the algal specimens collected from the various sampling sites in this work (**Fig. 1**) were compared with the microbiomes of *Ulva* species collected from Antarctica but maintained for approximately 30 years in the culture collection of the Alfred-Wegener Institute (AWI, Bremerhaven, Germany).

Most of the microbial community of *Ulva* specimens was taxonomically assigned to the phylum Proteobacteria, followed by Bacteroidota, as observed in many previous studies reporting that microbial communities associated with *Ulva* were taxonomically comparable (Friedrich 2012; Califano et al. 2020; Nguyen et al. 2023) but different from those associated with *Monostroma* species (**Fig. 3A**). The bacterial communities differed from those in coastal sediments and in periodically occurring water ponds (Krucon et al. 2021; Galván et al. 2023).

Moreover, the comparison of the native microbiomes with the mix of 443 isolated and cultivable bacteria, which constituted only a tiny percentage of the total diversity of the microbiome in general (Fig. 3A), confirmed the prevalence of easily cultivable Proteobacteria (Grueneberg et al. 2016).

Interestingly, the native microbiome contained diverse families (> 3%) within Bacteroidota (**Fig. 3B**), including Flavobacteriaceae, some genera of which, such as *Maribacter* and *Zobellia*, can release the morphogen thallusin (Matsuo et al. 2005; Weiss et al. 2017). Among them, the abundance of Flavobacteriaceae was greater in the microbiome of *Ulva* samples collected from sites P2 and P4 (UPC-100-P4 (18.5%), UPC-108-P4 (30.5%) and UPC-109-P4 (37.1%)) than in the microbiome of the samples of *Monostroma* spp. attached to the rocks. (**Fig. 3B, C**). In addition to the *Ulva* microbiome, the *Porphyra* microbiome also had a high abundance of Flavobacteriaceae. Therefore, *Ulva* and *Porphyra* were selected for more detailed microbiome analysis of Flavobacteriaceae.

In total, 22 different genera were assigned to Flavobacteriaceae (**Fig. 3C**), which were negligible in the free-floating *Monostroma* compared with *Ulva* or *Porphyra* (**Fig. 3B**). The abundances of Flavobacteriaceae were greater in sessile specimens, independent of the macroalga taxon, than in free-floating *Monostroma* species. In future research, the reasons for the dominance of Cyanophyceae and the absence of Bacteroidota on the surface of free-floating *Monostroma* compared to sessile species should be investigated (**Fig. 3A**).

The associated microbiome of *Ulva* isolate UPC-109-P2 included various genera of Flavobacteriaceae, such as *Algitalea* (10.8%), *Polaribacter* (7.6%), *Dokdonia* (5.1%), *Pibocella* (5.3%), and *Cellulophaga* (4.9%), as well as additional genera such as *Maribacter*, with abundances less than 1% (**Fig. 3C**). As *Maribacter* and related genera are essential for *Ulva* growth and morphogenesis, they must be considered part of the core microbiome of *Ulva* (**Fig. 3C**) (Weiss et al. 2017; Ulrich et al. 2022). For example, the genera *Pibocella* and *Maribacter* were abundant in *Ulva* samples, regardless of whether they were freshly isolated (UPC isolates) or cultivated (AWI isolates). *Maribacter* and *Pibocella* were dominant in many samples (UPC-108, PPC-10-P0, or 1002-AWI) that attracted our interest and to a lesser extent in *Monostroma* (MPC-90-PS).

Guided by the microbiome data, we selected these algae to isolate bacteria, which were transferred from the algal surface on agar plates by swabbing. Indeed, microbiome analysis of all mixed cultivable bacteria sampled from the Antarctic revealed successful isolation of Flavobacteriaceae. We hypothesized that some isolates could phenocopy *Maribacter* sp. and release AGMPFs such as thallusin. Changes in the microbiome associated with a specific trait, such as thallusin production by *Maribacter* spp., suggest the presence of a cold-adapted "core microbiome" that could promote *Ulva* growth and morphogenesis.

More details about the *Ulva* core bacterial communities are highlighted by recent studies showing that bacteria that share a symbiotic relationship with *Ulva* vary along an environmental salinity gradient (van der Loos et al. 2022) or in algal aquacultures (Califano et al. 2020). Significantly, changes in the composition of the microbial communities in the Antarctic Ocean can be driven by only a single strong temperature anomaly (Ilicic et al. 2023). However, the "core microbiome", a common set of bacteria or genes among a majority or vast majority of host specimens within a particular habitat (Turnbaugh et al., 2007), is essential for bacterial-macroalgal interactions. The core microbiome is the set of bacteria that play a vital role in ecosystem functioning or substantially impact host fitness and flexibility (Shade and Handelsman 2012). Shared functional genes rather than taxonomic entities are considered the key criteria underlying the structure of the "functional core" (Burke et al. 2011; Risely 2020). For instance, species of the genera *Sulfitobacter* and *Paracoccus* are known as *Roseovarius*-equivalent species with the ability to produce unknown AGMPFs (cell division-promoting factors) for supporting *Ulva* development (Ghaderiardakani et al. 2017). In contrast, the genus *Maribacter* cannot be simply replaced (Weiss et al. 2017).

In conclusion, the explorative microbiome study revealed the presence of bacterial clades that are crucial for the production of thallusin (**Fig. 3C**). We thus aimed to isolate strains under cold stress that can serve as phenocopies exhibiting the traits of the isolated *Maribacter* sp. MS6. The macroalgae with the highest abundance of Flavobacteriaceae were thus selected. The omnipresent and widespread activity of the unknown *Roseobacter* factor was investigated in parallel.

### 3.3 Bioassay-guided classification of cold-adapted bacteria based on their induction of morphogenesis in *Ulva mutabilis* Føyn

Bacterial isolates (*n* = 443) were successfully isolated, cultivated, and screened for the ability to induce cell division and differentiation to establish a novel tripartite community of *Ulva* with cold-adapted bacteria. As axenic cultures of the cold-adapted strain UPC-108 were unavailable, axenic cultures of *U. mutabilis* were used under standardized conditions (Grueneberg et al. 2016). Phenotypic analysis of *U. mutabilis* in the presence of cold-adapted bacteria was carried out three weeks after inoculation. The following categories (**I-V**) of bacterium-dependent algal phenotypes, which were originally defined by Grueneberg et al. (2016), were observed (see **Fig. 4A**):

**(I)** Bacteria showed “no morphogenetic activity,” and the algae developed a typical callus-like morphology with a lack of adherence and were covered with bubble-like protrusions due to distortions in the exterior cell wall (yellow circle).
**(II)** Bacterium-induced cell division was observed, such as with the reference strain *Roseovarius* sp. MS2 (magenta circle).
**(III)** Bacteria, like the reference strain *Maribacter* sp MS6, elicited cell differentiation (red circle).
**(IV)** Bacteria showed "algicidal activity" (black circle).
**(V)** Bacteria induced a "new morphotype," which was characterized by a flat and incomplete thallus (blue circle). Strains of categories II and III are the most intriguing in this study, as they are candidate strains for mitigating cold stress via bacterial–algal interactions.

Most of these bacterial strains (*n* = 326) had no morphogenetic activity (category I), and only 40 species could partially promote growth, but they did not solely elicit a complete morphotype (category II). Forty-seven isolates induced rhizoid and cell wall formation, potentially due to the release of thallusin (category III). In contrast, 12 isolates adversely affected *Ulva* germination (category IV), and 18 isolates induced the rarely observed flattened morphotype (category V) (**Fig. 4B**). In general, bacteria that phenocopied *Maribacter* sp. MS6 inhabited all the algal surfaces (**Table 2**). The *Ulva* bioassay-guided approach is the first step in obtaining the initial indication of bacteria-dependent morphogenesis in *Ulva*. However, this method can serve as a foundation for further studies investigating the underlying traits of bacteria involved in *Ulva* development in a changing environment and their strategies for adapting to stress (Ghaderiardakani et al. 2022).

**Table 2:** Overview of the determined morphogenetic activities of the isolated bacteria.

| **Origin** | **Axenic morphotype** | **Phenocopy of *Roseovarius* sp. MS2** | **Phenocopy of *Maribacter* sp. MS6** | **New flattened morphotype** | **Algicidal mode of action** |
| --- | --- | --- | --- | --- | --- |
| *Ulva* spp. | 115 | 13 | 13 | 5 | 4 |
| *Monostroma* spp. | 131 | 16 | 30 | 11 | 7 |
| *Porphyra* spp. | 26 | 5 | 4 | 0 | 1 |
| Sea water | 15 | 4 | 0 | 2 | 1 |
| **Total** | **327** | **38** | **47** | **18** | **13** |

### 3.4 Molecular identification for morphotype-based selection of the isolated bacteria

Fifty out of the 443 isolated bacteria were selected, depending on the morphotypes that they induced, for 16S rRNA-based phylogenetic identification, as determined by the *Ulva* bioassays. *Roseovarius* sp. MS2, *Maribacter* sp. MS6 and *Maribacter antarcticus* DSM 21422 were used as internal controls during sequencing.

Bacterial identification revealed that the isolated bacteria were dominated by *Pseudoalteromonas* spp., which does not confer morphogenetic activity (**Fig. 4C**). The isolates with the ability to induce cell differentiation and rhizoid formation (category III) were identified as *Lutibacter* sp. (BPC-D3) and *Winogradskyella s*p. (BPC-B5). Importantly, the isolate BPC-D8 neighbored *Maribacter* sp. MS6 and *Pibocella* sp., which are candidates for thallusin production. In contrast, as previously shown, the isolated *Algibacter* species (BPC-H2) were not morphogenetically active (Weiss et al. 2017).

Furthermore, the isolates BPC-C4 and BPC-B10, closely related to *Sulfitobacter brevis* DSM 11443 and *Paracoccus aestuarii* DSM 19484, induced cell division and growth (category II) (**Fig. 4C**). These yet-unknown AGMPFs are produced by a range of bacteria, such as *Roseovarius* sp. strain MS2, *Sulfitobacter* sp. strain MS3, *Halomonas* sp. strain MS1 and *Paracoccus* sp. (Grueneberg et al. 2016; Ghaderiardakani et al. 2017).

In summary, *U. mutabilis* recovered the typical morphotype in the presence of both newly isolated AGMPF-producing bacteria, BPC-D8 (*Maribacter* sp. MS6 equivalent) and BPC-C4 (*Roseovarius* sp. MS2 equivalent), collected from Antarctica (**Fig. 5**).

### 3.5 Whole-genome sequence of the novel cold-adapted thallusin-producing bacteria

The precise assignment of the bacterial isolate BPC-D8 to a species was achieved using PacBio sequencing technology. The draft genome of isolate BPC-D8 was assembled into one contig (4.7 Mbp), with a G+C content of 34.63% and an N_50_ value of 4783464 bp (**Table 3**, NCBI accession number: CP128187). Initial 16S rRNA gene-based sequencing indicated that the isolate was most closely related to *Maribacter aquivivus* and *Pibocella ponti*. Indeed, recent improvements in genomics and taxonomic assignments, such as the application of comparisons and standardized taxonomic classifications based on the ANI using GTDB-Tk (Chaumeil et al., 2020), have further demonstrated that the isolate is more closely affiliated with the suggested type strain *Maribacter aquivivus* DSM 16478 (**Fig. 6**). According to GTDB-Tk-v1.7.0 analysis, the ANI between BPC-D8 and *M. aquivivus* (GCA_900142175.1) was 90.55%, based on an aligned fraction of 0.82 (proportion of regions displaying significant similarity).

**Table 3:** Key parameters of the genome assembly of the strain BPC-D8.

| **Parameter** |  |
| --- | --- |
| **Genome size** | 4,783,464 bp |
| **GC content** | 34.6 |
| **Genome coverage** | 439 × |
| **Sequencing Technology** | PacBio |
| **Genes (total)** | 4,164 |
| **CDSs (total)** | 4,111 |
| **CDSs (coding)** | 4,099 |
| **Genes (RNA)** | 53 |
| **rRNAs (5S, 16S, 23S)** | 3, 3, 3 |
| **tRNAs** | 40 |
| **ncRNAs** | 4 |

The flavobacterial genus *Pibocella* sp. was first described by Nedashkovskaya et al. (2005) as a marine, heterotrophic, gram-negative, aerobic, yellow-pigmented, motile-by-gliding bacterium that was isolated from the green alga *Acrosiphonia sonderi* collected from the Sea of Japan. As determined by 16S rRNA gene sequence analysis, the new bacteria were assigned to the family Flavobacteriaceae, and the similarity values of the new isolate and its close relatives *Maribacter aquivivus*, *Maribacter orientalis*, and *Maribacter ulvicola* were 95.8%, 95.7% and 95.0%, respectively. Due to the low levels of sequence similarity between this isolate and the other known members of the Flavobacteriaceae family, *Pibocella ponti* gen. nov., sp. nov. was introduced (Nedashkovskaya et al. 2005). However, the taxonomic classification remains controversial, particularly at the genus and species levels (Bowman 2017; García-López et al. 2019). Although an inclusive taxonomic analysis on the classification of bacteria phylogenetically affiliated with the phylum Bacteroidota using a comprehensive set of aligned full-length 16S rRNA gene sequences nested *Pibocella* sp. within the *Maribacter* clade, a taxonomic revision for this genus using cutting-edge advanced genome sequences has been suggested (García-López et al. 2019). In summary, our whole-genome analysis suggested the classification of BPC-D8 into the *Maribacter* clade (**Fig. 6**), and the strain was thus named *Maribacter* sp. strain BPC-D8.

### 3.6. Growth of AGMPF-releasing bacteria and determination of thallus growth under cold conditions

Symbiotic bacteria such as *Roseovarius* sp. MS2 and *Maribacter* sp. MS6 release morphogenetic AGMPFs that facilitate proper morphogenesis of *Ulva* (Kessler et al. 2018). These bacterial metabolites are essential and may improve algal performance under stress. As a result, we propose that AGMPF and thallusin production could be important examples of a holobiont environmental stress response (**Fig. 7**). The investigated isolates *Maribacter* sp. MS6 and BPC-D8 were grown at low temperatures. Presumably, because of cold stress, the growth of the explored strains varied greatly (**Fig. 7A**). The growth of *Maribacter* sp. BPC-D8 began 12 days after inoculation and reached the stationary phase (OD_620_ = 0.24) after 24 days, while the growth of *Maribacter* sp. MS6 began after 28 days and reached the stationary phase (OD_620_ = 0.2) after 37 days. The growth rates (μ) during the exponential phase of these two bacteria were comparable at 2 °C (for *Maribacter* sp. MS6: μ = 0.01 ± 0.0002 h^−1^; for *Maribacter* sp. BPC-D8: μ = 0.009 ± 0.001 h^−1^) and approximately 10 times lower than that at 20 ± 1 °C (Ulrich et al. 2022). Interestingly, the temperate-adapted strain *Roseovarius* sp. MS2 did not grow at 2 °C (**Fig. 7B**); thus, it cannot provide the essential AGMPFs for *Ulva* growth.

**Figure 7.**
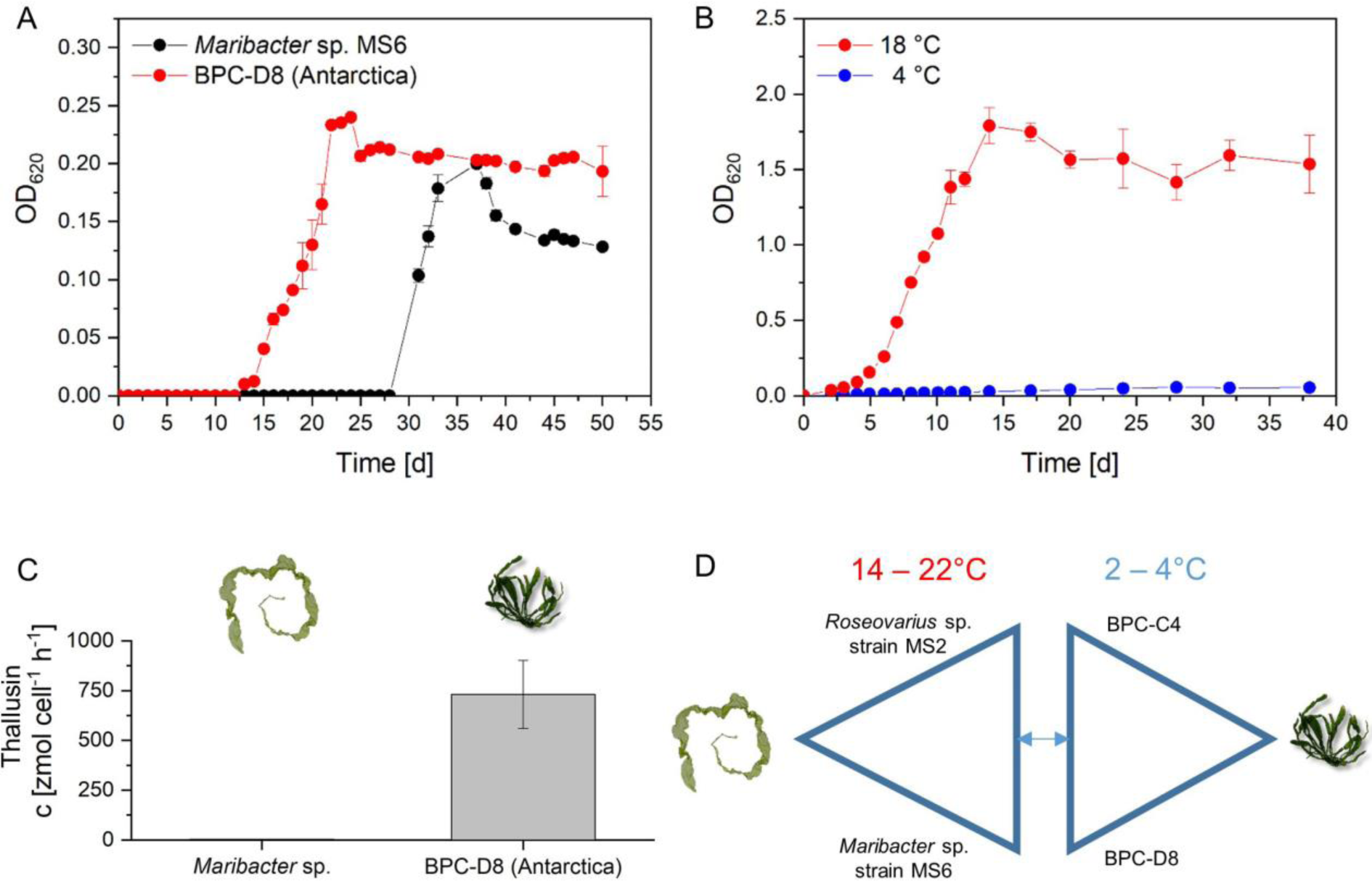
Growth of and thallusin production by bacteria that promoted algal growth and morphogenesis. (**A**) *Maribacter* sp. MS6 and the isolated *Maribacter* BPC-D8 strains were cultivated in HaHa_100 growth medium for seven weeks and maintained under cold conditions (2–4 °C). (**B**) *Roseovarius* sp. MS2 did not grow at 2 °C, but the isolated Sulfitobacter strain BPC-C4 did grow at this temperature. Both strains were cultivated in marine broth. (**C**) Although the temperate-adapted *Maribacter* sp. MS6 grew at 2–4 °C after the lag phase, the thallusin production rate was not significantly different. A small number of bacterial cells of the BPC-D8 isolate produced thalli at a high rate. (**D**) The proposed tripartite model system with *Ulva* and two essential algal growth and morphogenesis-promoting bacteria, *Maribacter* BPC-D8 and *Sulfitobacter* BPC-C4, was collected on King George Island (Potter Cove) in Antarctica.

Using mass spectrometry coupled with liquid chromatography, the effective concentration of thallusin was quantified in the culture medium of cold-adapted and temperate *Maribacter* strains at 2 °C. Samples were collected early in the exponential growth phase (OD_620_ = 0.05). Due to the increase in thallusin concentration with cell density and accumulation in the medium along with bacterial growth, the thallusin concentration was normalized to cell number and time (0.7 amol cell^-1^ h^-1^) (**Fig. 7C**). Considering the high biological activity of thallusin (Ulrich et al. 2022), the fast production of local high concentrations of thallusin, which was provided by the cold-adapted BPC-D8 and not by the warm-adapted strain (**Fig. 7C**), is more ecologically relevant, for example, in biofilms on the algal surface (Wahl et al. 2012) than in the culture medium.

In summary, the Mediterranean bacteria either did not grow at 2 °C or showed an extended lag phase after the temperature shift along with a decrease in thallusin production, which did not result in the initiation of germination of the released *Ulva* gametes or zoids. In future studies, the hypothesis of whether bacteria can support the transition of *Ulva* from a warm-temperate zone to an Antarctic zone and vice versa can be tested (**Fig. 7D**).

### 3.7. Limitations of the study

While our work focused on the identification of AGMPF-producing bacteria, seasonal studies will contribute to the understanding of changes in the dynamics of the microbiome and the release of growth-promoting factors in a changing environment. Thallusin-producing bacteria were found in red and green algae, which suggests intriguing eco-physiological functions. However, functional studies with *Porphyra* were not performed because axenic cultures needed to be established first (Califano and Wichard 2018). Furthermore, the regulation of gametogenesis in isolated Antarctic *Ulva* species should be intensively studied as a prerequisite for achieving axenic cultures of *Ulva* sp. strain BPC-108 under controlled conditions and propagation of gametophytes similar to those of *U. mutabilis*. Finally, our study did not participate in the controversial taxonomic reclassification of *Monostroma* (Cui et al. 2022). Therefore, further phylogenetic studies are needed.

## 4. Conclusions

Our work contributes to key questions related to stress ecology in marine ecosystems, which might also be relevant to the environmental impacts of climate change. Unlike the temperate *U. mutabilis,* the Antarctic *Ulva* strains grew and released gametes, which developed into gametophytes at 2 °C. The regulation of sporulation needs to be further investigated to manage the Antarctic *Ulva* life cycle according to established protocols (Stratmann et al. 1996). Interestingly, characteristic bacterial communities can contribute to the adaptation mechanisms of *Ulva* and facilitate its performance by exposing it to environmental stresses. We isolated and identified *Ulva*-associated and AGMPF-providing bacteria within the algal growth-promoting polar microbiome. The ability of Antarctic bacteria to release growth-promoting compounds under cold conditions was demonstrated by detecting and quantifying thallusin in the culture medium. The obtained results will serve as a foundation for future in-depth studies that integrate field and experimental work and focus more on causal relationships between bacteria and algal hosts in response to cold or combined stressors, which are more typical in the natural environment than single stressors. Exploring the functional selection of bacterial communities under different (single or combined) stresses could clarify whether an *Ulva*-associated set of core functional traits for adaptation exists and whether the bacterial taxonomic core could be connected to such functional arrangements. An enhanced understanding of stress mitigation strategies and adaptation mechanisms will help to reduce the impact of these stresses on biodiversity or ecosystem elements.

## Data availability

The sequencing reads and assemblies for this whole-genome shotgun project are available in the National Center for Biotechnology Information repository under the BioProject accession number PRJNA828511. The version described in this paper is the first version (NCBI accession number: CP128187). The complete data set of 16S rDNA sequences from the isolated bacteria and the surface microbiome analyses have been deposited at NCBI with the Accession numbers OR957431-OR957479 and SAMN39202330-SAMN39202384 under the same BioProject. Sequence data of the *tufA* gene have the accession numbers PP091296-PP091308. The provided Sequence IDs are identical to those used in this publication.

## Acknowledgments

This work was funded by the Deutsche Forschungsgemeinschaft (DFG, German Research Foundation) in the framework of the priority program (SPP 1158) "Antarctic Research with comparative investigations in Arctic ice areas" (424256657, FG, and TW) and of SFB 1127/2 “ChemBioSys” (239748522, TW). The study received financial support from the Ministry for Economics, Sciences and Digital Society of Thuringia (TMWWDG) under the framework of the Landesprogramm ProDigital (DigLeben-5575/10-9). JFU was funded by a doctoral scholarship from the Konrad-Adenauer-Stiftung. MQ was funded by ANPCyT-DNA PICT 2691 and 0501. Finally, we thank the Carlini Station crews for their support, logistics, and hospitality. We also appreciate the support of the German sampling team: Hans-Peter Grossart, Ulf Karsten, Alexandra Livenets, and Jonas Zimmermann. We thank Georg Pohnert (Friedrich-Schiller-Universität Jena) for his great support in Jena.

## Author contribution statement

**Fatemeh Ghaderiardakani**: Formal analysis, Investigation, Methodology, Validation, Visualization, Writing–original draft, Writing–review & editing. **Johann F. Ulrich**: Investigation Visualization, Writing–review & editing. **Emanuel Barth**: Formal analysis Methodology. **Maria Liliana Quartino**: Investigation, Project administration, Supervision, Writing– review & editing. **Thomas Wichard**: Conceptualization, Data curation, Funding acquisition, Methodology, Project administration, Supervision, Validation, Writing–original draft, Writing–review & editing

