## Supporting Information for "Algal growth and morphogenesis-promoting factors released by cold-adapted bacteria contribute to the resilience and morphogenesis of the seaweed *Ulva* (Chlorophyta) in Antarctica (Potter Cove)"

Table S1: Isolated green and red macroalgae from seawater

| **Sequence ID/ Starin_ID** | Accession number | **Collection date** | **Collected by** | **Country** | Culture **collection** | **Note** |
|---|---|---|---|---|---|---|
| **UPC-106** | PP091296 | Feb 2020 | F. Ghaderiardakani, T. Wichard | Antarctica: Isla 25 de Mayo/ King George Island, Potter Cove | --- | |
| **UPC-107** | PP091297 | Feb 2020 | F. Ghaderiardakani, T. Wichard | Antarctica: Isla 25 de Mayo/ King George Island, Potter Cove | --- | |
| **UPC-108** | PP091298 | Feb 2020 | F. Ghaderiardakani, T. Wichard | Antarctica: Isla 25 de Mayo/ King George Island, Potter Cove | --- | |
| **UPC-109** | PP091299 | Feb 2020 | F. Ghaderiardakani, T. Wichard | Antarctica: Isla 25 de Mayo/ King George Island, Potter Cove | --- | |
| **MPC-51** | PP091300 | Feb 2020 | F. Ghaderiardakani, T. Wichard | Antarctica: Isla 25 de Mayo/ King George Island, Potter Cove | --- | |
| **MPC-54** | PP091301 | Feb 2020 | F. Ghaderiardakani, T. Wichard | Antarctica: Isla 25 de Mayo/ King George Island, Potter Cove | --- | |
| **MPC-63** | PP091302 | Feb 2020 | F. Ghaderiardakani, T. Wichard | Antarctica: Isla 25 de Mayo/ King George Island, Potter Cove | --- | |
| **MPC-90** | PP091303 | Feb 2020 | F. Ghaderiardakani, T. Wichard | Antarctica: Isla 25 de Mayo/ King George Island, Potter Cove | --- | |
| **PPC-66** | PP091304 | Feb 2020 | F. Ghaderiardakani, T. Wichard | Antarctica: Isla 25 de Mayo/ King George Island, Potter Cove | --- | |
| **PPC-73** | PP091305 | Feb 2020 | F. Ghaderiardakani, T. Wichard | Antarctica: Isla 25 de Mayo/ King George Island, Potter Cove | --- | |
| **AWI_1002** | PP091306 | 1986 | C. Wiencke | Antarctica: Isla 25 de Mayo/ King George Island, Jubany station | Alfred Wegener Institute, Bremerhaven, Germany | |
| **AWI_1158** | PP091307 | 1994 | C. Wiencke | Antarctica: Isla 25 de Mayo/ King George Island, Jubany station | Alfred Wegener Institute, Bremerhaven, Germany | |
| **AWI_1159** | PP091308 | 1994 | C. Wiencke | Antarctica: Isla 25 de Mayo/ King George Island, Jubany station | Alfred Wegener Institute, Bremerhaven, Germany | |
| **FSU-UM5-1** | PP091309 | 1952 | B. Føyn | Portugal: Ria Formosa | Friedrich Schiller University Jena, Germany | Morphotype, slender |

Table S2: Samples of the microbiome analysis

| NCBI accession | Sample name | Host | Collection date | Geographic location | Latitude_and_longitude | Sample type |
|---|---|---|---|---|---|---|
| **SAMN39202330** | PPC-10-P0-1 | *Porphyra* | 2020-02 | Antarctica: Isla 25 de Mayo/ King George Island, Potter Cove | not collected | replicate 1 |
| **SAMN39202331** | PPC-10-P0-2 | *Porphyra* | 2020-02 | Antarctica: Isla 25 de Mayo/ King George Island, Potter Cove | not collected | replicate 2 |
| **SAMN39202332** | PPC-10-P0-3 | *Porphyra* | 2020-02 | Antarctica: Isla 25 de Mayo/ King George Island, Potter Cove | not collected | replicate 3 |
| **SAMN39202333** | MPC-27-PDP-1 | *Monostroma* | 2020-02 | Antarctica: Isla 25 de Mayo/ King George Island, Potter Cove | 62.238 S 58.705 W | replicate 1 |
| **SAMN39202334** | MPC-27-PDP-2 | *Monostroma* | 2020-02 | Antarctica: Isla 25 de Mayo/ King George Island, Potter Cove | 62.238 S 58.705 W | replicate 2 |
| **SAMN39202335** | MPC-27-PDP-3 | *Monostroma* | 2020-02 | Antarctica: Isla 25 de Mayo/ King George Island, Potter Cove | 62.238 S 58.705 W | replicate 3 |
| **SAMN39202336** | MPC-28-PDP-1 | *Monostroma* | 2020-02 | Antarctica: Isla 25 de Mayo/ King George Island, Potter Cove | 62.238 S 58.705 W | replicate 1 |
| **SAMN39202337** | MPC-28-PDP-2 | *Monostroma* | 2020-02 | Antarctica: Isla 25 de Mayo/ King George Island, Potter Cove | 62.238 S 58.705 W | replicate 2 |
| **SAMN39202338** | MPC-28-PDP-3 | *Monostroma* | 2020-02 | Antarctica: Isla 25 de Mayo/ King George Island, Potter Cove | 62.238 S 58.705 W | replicate 3 |
| **SAMN39202339** | MPC-29-A2-1 | *Monostroma* | 2020-02 | Antarctica: Isla 25 de Mayo/ King George Island, Potter Cove | 62.234 S 58.668 W | replicate 1 |
| **SAMN39202340** | MPC-29-A2-2 | *Monostroma* | 2020-02 | Antarctica: Isla 25 de Mayo/ King George Island, Potter Cove | 62.234 S 58.668 W | replicate 2 |
| **SAMN39202341** | MPC-29-A2-3 | *Monostroma* | 2020-02 | Antarctica: Isla 25 de Mayo/ King George Island, Potter Cove | 62.234 S 58.668 W | replicate 3 |
| **SAMN39202342** | MPC-38-A2-1 | *Monostroma* | 2020-02 | Antarctica: Isla 25 de Mayo/ King George Island, Potter Cove | 62.234 S 58.668 W | replicate 1 |
| **SAMN39202343** | MPC-38-A2-2 | *Monostroma* | 2020-02 | Antarctica: Isla 25 de Mayo/ King George Island, Potter Cove | 62.234 S 58.668 W | replicate 2 |
| **SAMN39202344** | MPC-38-A2-3 | *Monostroma* | 2020-02 | Antarctica: Isla 25 de Mayo/ King George Island, Potter Cove | 62.234 S 58.668 W | replicate 3 |
| **SAMN39202345** | MPC-57-A4-1 | *Monostroma* | 2020-02 | Antarctica: Isla 25 de Mayo/ King George Island, Potter Cove | 62.228 S 58.664 W | replicate 1 |
| **SAMN39202346** | MPC-57-A4-2 | *Monostroma* | 2020-02 | Antarctica: Isla 25 de Mayo/ King George Island, Potter Cove | 62.228 S 58.664 W | replicate 2 |
| **SAMN39202347** | MPC-57-A4-3 | *Monostroma* | 2020-02 | Antarctica: Isla 25 de Mayo/ King George Island, Potter Cove | 62.228 S 58.664 W | replicate 3 |
| **SAMN39202348** | MPC-51-PE-1 | *Monostroma* | 2020-02 | Antarctica: Isla 25 de Mayo/ King George Island, Potter Cove | 62.237 S 58.678 W | replicate 1 |
| **SAMN39202349** | MPC-51-PE-2 | *Monostroma* | 2020-02 | Antarctica: Isla 25 de Mayo/ King George Island, Potter Cove | 62.237 S 58.678 W | replicate 2 |
| **SAMN39202350** | MPC-51-PE-3 | *Monostroma* | 2020-02 | Antarctica: Isla 25 de Mayo/ King George Island, Potter Cove | 62.237 S 58.678 W | replicate 3 |
| **SAMN39202351** | MPC-64-PE-1 | *Monostroma* | 2020-02 | Antarctica: Isla 25 de Mayo/ King George Island, Potter Cove | 62.237 S 58.678 W | replicate 1 |
| **SAMN39202352** | MPC-64-PE-2 | *Monostroma* | 2020-02 | Antarctica: Isla 25 de Mayo/ King George Island, Potter Cove | 62.237 S 58.678 W | replicate 2 |
| **SAMN39202353** | MPC-64-PE-3 | *Monostroma* | 2020-02 | Antarctica: Isla 25 de Mayo/ King George Island, Potter Cove | 62.237 S 58.678 W | replicate 3 |
| **SAMN39202354** | MPC-90-PS-1 | *Monostroma* | 2020-02 | Antarctica: Isla 25 de Mayo/ King George Island, Potter Cove | 62.262 S 58.618 W | replicate 1 |

| | | | | | | |
|---|---|---|---|---|---|---|
| **SAMN39202355** | MPC-90-PS-2 | *Monostroma* | 2020-02 | Antarctica: Isla 25 de Mayo/ King George Island, Potter Cove | 62.262 S 58.618 W | replicate 2 |
| **SAMN39202356** | MPC-90-PS-3 | *Monostroma* | 2020-02 | Antarctica: Isla 25 de Mayo/ King George Island, Potter Cove | 62.262 S 58.618 W | replicate 3 |
| **SAMN39202357** | UPC-100-P4-1 | *Ulva* | 2020-02 | Antarctica: Isla 25 de Mayo/ King George Island, Potter Cove | 62.255 S 58.665 W | replicate 1 |
| **SAMN39202358** | UPC-100-P4-2 | *Ulva* | 2020-02 | Antarctica: Isla 25 de Mayo/ King George Island, Potter Cove | 62.255 S 58.665 W | replicate 2 |
| **SAMN39202359** | UPC-100-P4-3 | *Ulva* | 2020-02 | Antarctica: Isla 25 de Mayo/ King George Island, Potter Cove | 62.255 S 58.665 W | replicate 3 |
| **SAMN39202360** | UPC-108-P2-1 | *Ulva* | 2020-02 | Antarctica: Isla 25 de Mayo/ King George Island, Potter Cove | 62.251 S 58.675 W | replicate 1 |
| **SAMN39202361** | UPC-108-P2-2 | *Ulva* | 2020-02 | Antarctica: Isla 25 de Mayo/ King George Island, Potter Cove | 62.251 S 58.675 W | replicate 2 |
| **SAMN39202362** | UPC-108-P2-3 | *Ulva* | 2020-02 | Antarctica: Isla 25 de Mayo/ King George Island, Potter Cove | 62.251 S 58.675 W | replicate 3 |
| **SAMN39202363** | UPC-109-P2-1 | *Ulva* | 2020-02 | Antarctica: Isla 25 de Mayo/ King George Island, Potter Cove | 62.251 S 58.675 W | replicate 1 |
| **SAMN39202364** | UPC-109-P2-2 | *Ulva* | 2020-02 | Antarctica: Isla 25 de Mayo/ King George Island, Potter Cove | 62.251 S 58.675 W | replicate 2 |
| **SAMN39202365** | UPC-109-P2-3 | *Ulva* | 2020-02 | Antarctica: Isla 25 de Mayo/ King George Island, Potter Cove | 62.251 S 58.675 W | replicate 3 |
| **SAMN39202366** | UPC-114-P2-1 | *Ulva* | 2020-02 | Antarctica: Isla 25 de Mayo/ King George Island, Potter Cove | 62.251 S 58.675 W | replicate 1 |
| **SAMN39202367** | UPC-114-P2-2 | *Ulva* | 2020-02 | Antarctica: Isla 25 de Mayo/ King George Island, Potter Cove | 62.251 S 58.675 W | replicate 2 |
| **SAMN39202368** | UPC-114-P2-3 | *Ulva* | 2020-02 | Antarctica: Isla 25 de Mayo/ King George Island, Potter Cove | 62.251 S 58.675 W | replicate 3 |
| **SAMN39202369** | UPC-115-P2-1 | *Ulva* | 2020-02 | Antarctica: Isla 25 de Mayo/ King George Island, Potter Cove | 62.251 S 58.675 W | replicate 1 |
| **SAMN39202370** | UPC-115-P2-2 | *Ulva* | 2020-02 | Antarctica: Isla 25 de Mayo/ King George Island, Potter Cove | 62.251 S 58.675 W | replicate 2 |
| **SAMN39202371** | UPC-115-P2-3 | *Ulva* | 2020-02 | Antarctica: Isla 25 de Mayo/ King George Island, Potter Cove | 62.251 S 58.675 W | replicate 3 |
| **SAMN39202372** | UPC-116-P2-1 | *Ulva* | 2020-02 | Antarctica: Isla 25 de Mayo/ King George Island, Potter Cove | 62.251 S 58.675 W | replicate 1 |
| **SAMN39202373** | UPC-116-P2-2 | *Ulva* | 2020-02 | Antarctica: Isla 25 de Mayo/ King George Island, Potter Cove | 62.251 S 58.675 W | replicate 2 |
| **SAMN39202374** | UPC-116-P2-3 | *Ulva* | 2020-02 | Antarctica: Isla 25 de Mayo/ King George Island, Potter Cove | 62.251 S 58.675 W | replicate 3 |
| **SAMN39202375** | 1158-AWI-1 | *Ulva* | 2020-02 | Antarctica: Isla 25 de Mayo/ King George Island, Potter Cove, Jubany station | 62.233 S 58.667 W | replicate 1 |
| **SAMN39202376** | 1158-AWI-2 | *Ulva* | 2020-02 | Antarctica: Isla 25 de Mayo/ King George Island, Potter Cove, Jubany station | 62.233 S 58.667 W | replicate 2 |
| **SAMN39202377** | 1158-AWI-3 | *Ulva* | 2020-02 | Antarctica: Isla 25 de Mayo/ King George Island, Potter Cove, Jubany station | 62.233 S 58.667 W | replicate 3 |
| **SAMN39202378** | 1159-AWI-1 | *Ulva* | 2020-02 | Antarctica: Isla 25 de Mayo/ King George Island, Potter Cove, Jubany station | 62.233 S 58.667 W | replicate 1 |
| **SAMN39202379** | 1159-AWI-2 | *Ulva* | 2020-02 | Antarctica: Isla 25 de Mayo/ King George Island, Potter Cove, Jubany station | 62.233 S 58.667 W | replicate 2 |
| **SAMN39202380** | 1159-AWI-3 | *Ulva* | 2020-02 | Antarctica: Isla 25 de Mayo/ King George Island, Potter Cove, Jubany station | 62.233 S 58.667 W | replicate 3 |
| **SAMN39202381** | Mixed cultured bacteria for microbiome analysis 1 | macroalgae | 2020-02 | Antarctica: Isla 25 de Mayo/ King George Island, Potter Cove | not provided | replicate 1 |
| **SAMN39202382** | Mixed cultured bacteria for microbiome analysis 2 | macroalgae | 2020-02 | Antarctica: Isla 25 de Mayo/ King George Island, Potter Cove | not provided | replicate 2 |

| | | | | | | |
|---|---|---|---|---|---|---|
| **SAMN39202383** | Mixed cultured bacteria for microbiome analysis 3 | macroalgae | 2020-02 | Antarctica: Isla 25 de Mayo/ King George Island, Potter Cove | not provided | replicate 3 |
| **SAMN39202384** | Mixed cultured bacteria for microbiome analysis 4 | macroalgae | 2020-02 | Antarctica: Isla 25 de Mayo/ King George Island, Potter Cove | not provided | replicate 4 |

Table S3: Isolated Antarctic marine bacteria

| **Strain** | **Accession Number** | **Date** | **Sampling site** | **Source** | **Species** |
|---|---|---|---|---|---|
| **BPC-A5** | OR957431 | Feb 2020 | Antarctica: Isla 25 de Mayo/ King George Island, Potter Cove | Green macroalgae | *Monostroma* sp. |
| **BPC-A6** | OR957432 | Feb 2020 | Antarctica: Isla 25 de Mayo/ King George Island, Potter Cove | Green macroalgae | *Monostroma* sp. |
| **BPC-A7** | OR957433 | Feb 2020 | Antarctica: Isla 25 de Mayo/ King George Island, Potter Cove | Green macroalgae | *Monostroma* sp. |
| **BPC-A8** | OR957434 | Feb 2020 | Antarctica: Isla 25 de Mayo/ King George Island, Potter Cove | Green macroalgae | *Monostroma* sp. |
| **BPC-A9** | OR957435 | Feb 2020 | Antarctica: Isla 25 de Mayo/ King George Island, Potter Cove | Green macroalgae | *Monostroma* sp. |
| **BPC-A10** | OR957436 | Feb 2020 | Antarctica: Isla 25 de Mayo/ King George Island, Potter Cove | Sea water | |
| **BPC-B1** | OR957437 | Feb 2020 | Antarctica: Isla 25 de Mayo/ King George Island, Potter Cove | Red macroalgae | *Porphyra* sp. |
| **BPC-B4** | OR957438 | Feb 2020 | Antarctica: Isla 25 de Mayo/ King George Island, Potter Cove | Sea water | |
| **BPC-B5** | OR957439 | Feb 2020 | Antarctica: Isla 25 de Mayo/ King George Island, Potter Cove | Red macroalgae | *Porphyra* sp. |
| **BPC-B7** | OR957440 | Feb 2020 | Antarctica: Isla 25 de Mayo/ King George Island, Potter Cove | Sea water | |
| **BPC-B8** | OR957441 | Feb 2020 | Antarctica: Isla 25 de Mayo/ King George Island, Potter Cove | Red macroalgae | *Porphyra* sp. |
| **BPC-B10** | OR957442 | Feb 2020 | Antarctica: Isla 25 de Mayo/ King George Island, Potter Cove | Red macroalgae | *Porphyra* sp. |
| **BPC-B12** | OR957443 | Feb 2020 | Antarctica: Isla 25 de Mayo/ King George Island, Potter Cove | Red macroalgae | *Porphyra* sp. |
| **BPC-C1** | OR957444 | Feb 2020 | Antarctica: Isla 25 de Mayo/ King George Island, Potter Cove | Green macroalgae | *Monostroma* sp. |
| **BPC-C3** | OR957445 | Feb 2020 | Antarctica: Isla 25 de Mayo/ King George Island, Potter Cove | Green macroalgae | *Monostroma* sp. |
| **BPC-C4** | OR957446 | Feb 2020 | Antarctica: Isla 25 de Mayo/ King George Island, Potter Cove | Red and green macroalgae | *Ulva* sp. UPC-109 |
| **BPC-C5** | OR957447 | Feb 2020 | Antarctica: Isla 25 de Mayo/ King George Island, Potter Cove | Green macroalgae | *Monostroma* sp. |
| **BPC-C6** | OR957448 | Feb 2020 | Antarctica: Isla 25 de Mayo/ King George Island, Potter Cove | Sea water | |
| **BPC-C8** | OR957449 | Feb 2020 | Antarctica: Isla 25 de Mayo/ King George Island, Potter Cove | Green macroalgae | *Monostroma* sp. |
| **BPC-C10** | OR957450 | Feb 2020 | Antarctica: Isla 25 de Mayo/ King George Island, Potter Cove | Sea water | |
| **BPC-C11** | OR957451 | Feb 2020 | Antarctica: Isla 25 de Mayo/ King George Island, Potter Cove | Sea water | |

| | | | | | |
|---|---|---|---|---|---|
| **BPC-C12** | OR957452 | Feb 2020 | Antarctica: Isla 25 de Mayo/ King George Island, Potter Cove | Sea water | |
| **BPC-D1** | OR957453 | Feb 2020 | Antarctica: Isla 25 de Mayo/ King George Island, Potter Cove | Green macroalgae | *Monostroma* sp. |
| **BPC-D2** | OR957454 | Feb 2020 | Antarctica: Isla 25 de Mayo/ King George Island, Potter Cove | Sea water | |
| **BPC-D3** | OR957455 | Feb 2020 | Antarctica: Isla 25 de Mayo/ King George Island, Potter Cove | Green macroalgae | *Ulva* sp. UPC-109 |
| **BPC-D4** | OR957456 | Feb 2020 | Antarctica: Isla 25 de Mayo/ King George Island, Potter Cove | Red macroalgae | *Porphyra* sp. |
| **BPC-D8** | OR957457 | Feb 2020 | Antarctica: Isla 25 de Mayo/ King George Island, Potter Cove | Red and green macroalgae | *Ulva* sp. UPC-109 |
| **BPC-E1** | OR957458 | Feb 2020 | Antarctica: Isla 25 de Mayo/ King George Island, Potter Cove | Sea water | |
| **BPC-E2** | OR957459 | Feb 2020 | Antarctica: Isla 25 de Mayo/ King George Island, Potter Cove | Green macroalgae | *Ulva* sp. UPC-109 |
| **BPC-E3** | OR957460 | Feb 2020 | Antarctica: Isla 25 de Mayo/ King George Island, Potter Cove | Green macroalgae | *Ulva* sp. UPC-109 |
| **BPC-E5** | OR957461 | Feb 2020 | Antarctica: Isla 25 de Mayo/ King George Island, Potter Cove | Green macroalgae | *Monostroma* sp. |
| **BPC-F3** | OR957462 | Feb 2020 | Antarctica: Isla 25 de Mayo/ King George Island, Potter Cove | Red macroalgae | *Porphyra* sp. |
| **BPC-F4** | OR957463 | Feb 2020 | Antarctica: Isla 25 de Mayo/ King George Island, Potter Cove | Red macroalgae | *Porphyra* sp. |
| **BPC-F5** | OR957464 | Feb 2020 | Antarctica: Isla 25 de Mayo/ King George Island, Potter Cove | Sea water | |
| **BPC-F8** | OR957465 | Feb 2020 | Antarctica: Isla 25 de Mayo/ King George Island, Potter Cove | Green Macroalgae | *Monostroma* sp. |
| **BPC-F9** | OR957466 | Feb 2020 | Antarctica: Isla 25 de Mayo/ King George Island, Potter Cove | Red macroalgae | *Porphyra* sp. |
| **BPC-F11** | OR957467 | Feb 2020 | Antarctica: Isla 25 de Mayo/ King George Island, Potter Cove | Green macroalgae | *Monostroma* sp. |
| **BPC-G2** | OR957468 | Feb 2020 | Antarctica: Isla 25 de Mayo/ King George Island, Potter Cove | Green macroalgae | *Monostroma* sp. |
| **BPC-G3** | OR957469 | Feb 2020 | Antarctica: Isla 25 de Mayo/ King George Island, Potter Cove | Red macroalgae | *Porphyra* sp. |
| **BPC-G4** | OR957470 | Feb 2020 | Antarctica: Isla 25 de Mayo/ King George Island, Potter Cove | Green macroalgae | *Monostroma* sp. |
| **BPC-G6** | OR957471 | Feb 2020 | Antarctica: Isla 25 de Mayo/ King George Island, Potter Cove | Green macroalgae | *Monostroma* sp. |
| **BPC-G11** | OR957472 | Feb 2020 | Antarctica: Isla 25 de Mayo/ King George Island, Potter Cove | Red macroalgae | *Porphyra* sp. |
| **BPC-H1** | OR957473 | Feb 2020 | Antarctica: Isla 25 de Mayo/ King George Island, Potter Cove | Green macroalgae | *Monostroma* sp. |
| **BPC-H2** | OR957474 | Feb 2020 | Antarctica: Isla 25 de Mayo/ King George Island, Potter Cove | Sea water | |

| | | | | | |
|---|---|---|---|---|---|
| **BPC-H4** | OR957475 | Feb 2020 | Antarctica: Isla 25 de Mayo/ King George Island, Potter Cove | Red macroalgae | *Porphyra* sp. |
| **BPC-H5** | OR957476 | Feb 2020 | Antarctica: Isla 25 de Mayo/ King George Island, Potter Cove | Green macroalgae | *Ulva* sp. UPC-109 |
| **BPC-H6** | OR957477 | Feb 2020 | Antarctica: Isla 25 de Mayo/ King George Island, Potter Cove | Red macroalgae | *Porphyra* sp. |
| **BPC-H10** | OR957478 | Feb 2020 | Antarctica: Isla 25 de Mayo/ King George Island, Potter Cove | Sea water | |
| **BPC-H12** | OR957479 | Feb 2020 | Antarctica: Isla 25 de Mayo/ King George Island, Potter Cove | Sea water | |
